# Pleiotropy and facilitation of local adaptation in the silverleaf sunflower *Helianthus argophyllus*

**DOI:** 10.1101/2025.05.22.655459

**Authors:** Uzezi E. Okinedo, Brook T. Moyers

## Abstract

Local adaptation drives changes in population phenotypes that confer survival or increased reproduction in certain environments. Local adaptation may be hindered or facilitated by pleiotropy, which is the control of multiple traits by a single genetic locus. In this study, we explored the connections between pleiotropy and local adaptation in the Texas endemic silverleaf sunflower, *Helianthus argophyllus*. Populations of *H. argopyllus* exhibit a bimodal life history strategy, consisting of tall, late-flowering forms and short, early-flowering forms that occur in close geographic proximity. The expression of life history traits within *H. argophyllus* populations is linked to local adaptation and controlled by a highly pleiotropic locus. Still, we do not know how local adaptation and pleiotropy interact at the transcriptomic level. We identified putatively locally adapted genes using whole RNA sequencing data and two selection outlier approaches. We assessed transcriptomic pleiotropy by evaluating whether allelic variants within genes control the expression of other genes (an eQTL approach) and gene co-expression network connectivity. Our results show that candidate locally adaptive genes identified by both methods are enriched for eQTL loci. Candidate genes identified through associations with environmental variables exhibit modular expression and have significantly lower network connectivity than non-adapted genes. In contrast, candidate genes identified only by controlling for population structure have significantly higher network connectivity. Our results suggest that the role of pleiotropy in local adaptation, at least at the transcriptomic level, depends on the function of the specific loci under selection.

**Author Summary:** This study investigates how genes contribute to local adaptation in *Helianthus argophyllus*, a sunflower species with distinct ecotypes. Using transcriptomic data and two methods involving environmental association and population structure analysis, we found that adaptive genes can be categorized into two types: pleiotropic “hub” genes, which affect multiple traits, and modular genes, which affect fewer, more specific traits. Environment-associated genes tended to be modular, while population-structure-associated genes were more pleiotropic. These patterns support a dynamic model of adaptation, where pleiotropy is beneficial early in adaptation and modularity becomes increasingly crucial as populations refine their traits. The findings provide new insights into how gene network architecture influences evolutionary responses to environmental change, particularly in diverging populations, such as those of *H. argophyllus*.

## Introduction

Pleiotropy is an important concept in adaptive evolution. It describes how a single genetic locus influences multiple phenotypic traits [1,2]. It can arise from the effects of shared cis-regulatory elements on gene expression or from the joint influence of gene networks, forming variational modules [3]. These modules often represent gene co-expression networks, where connections between trait nodes and gene nodes indicate genetic influence on phenotype expression [3,4]. Understanding how pleiotropy shapes adaptation is critical for assessing whether adaptive loci function as regulatory hubs or if selection favors more specialized genetic architectures.

Evolutionary models, such as Fisher’s geometric model and the cost of pleiotropy theory, suggest that highly pleiotropic or large-effect loci may constrain adaptation by generating trade-offs that shift an organism from its fitness optimum [5,6]. Consequently, adaptation often proceeds through small-effect loci or modular subunits, which minimize detrimental consequences on an organism’s fitness. However, extensions of Fisher’s theory propose that pleiotropic loci can accelerate adaptation, mainly when populations are far from their fitness optimum, such as during habitat expansion or environmental shifts [1,6]. Experimental evidence supports this by showing that adaptive loci with larger effect sizes often correlate with increased distance from optimal fitness [4,7]. This raises the question of whether locally adaptive loci tend to be pleiotropic or more functionally specialized, a key focus of this study.

*Helianthus argophyllus* (Silverleaf sunflower) is an annual species in the Asteraceae family, endemic to Texas and parts of Florida, and grows in sandy soils across diverse habitats [8][9]. It shares morphological similarities with *Helianthus annuus* but is distinguishable by dense silvery-white pubescence on its leaves, stems, and phyllaries [10,11]. A key feature of *H. argophyllus* is its bimodal life history strategy, which has led to the emergence of two distinct ecotypes: a tall, late-flowering ecotype with low branching found in drier inland Texas and a short, early-flowering, highly branched ecotype, distributed on barrier islands along the Gulf of Mexico [12,13]. Despite their narrow geographical range, these ecotypes coexist within the same populations but differ in flowering phenology [13]. The tall, late-flowering ecotype blooms from late August to early September and is the primary ecotype in inland populations.

In contrast, the short, early-flowering ecotype blooms in June and is more common in coastal populations [9]. This bimodal flowering strategy is controlled by HaFT1, a FLOWERING LOCUS T homolog introgressed from *H. annuus*. HaFT1 is located in a 10Mb low-recombination region on chromosome 6 of *H. annuus*, and likely in the same position in *H. argophyllus,* although this chromosome has a reciprocal translocation with *H. annuus* chromosome 15. The short, early-flowering ecotype possesses one or both copies of HaFT1, either homozygous or heterozygous for its presence, representing the ancestral allele. In contrast, the tall, late-flowering ecotype lacks HaFT1 entirely (homozygous for its absence, the derived allele) [8,14].

Some studies suggest that adaptive loci are modular and minimize pleiotropic constraints, while others argue that pleiotropy is a key feature of adaptive evolution, allowing correlated trait changes [4,15–17]. However, few studies have directly examined whether adaptive loci in naturally diverging populations tend to be pleiotropic or functionally specialized. Understanding this relationship is critical for determining whether selection favors genetic architectures that minimize trade-offs or integrate multiple adaptive changes through pleiotropic loci [18].

*Helianthus argophyllus* provides an ideal system for studying pleiotropy in adaptation because it exhibits clear ecotypic divergence with a known genetic basis. The HaFT1 locus is strongly associated with flowering time, which correlates with fitness-related traits such as seed size, seed mass, and plant height [13,14,19–21]. If pleiotropic loci drive adaptation in this system, we would expect adaptive genes to function as regulatory hubs influencing multiple traits. Alternatively, if selection favors modularity, adaptive loci may exhibit low pleiotropy, with distinct genes controlling different adaptive traits.

In this study, we apply a transcriptomic and network-based approach to investigate the role of pleiotropy in the local adaptation of *H. argophyllus*. To estimate genetic diversity indices, we sequence the transcriptomes of individuals representing both tall, late-flowering, and short, early-flowering ecotypes. We identify adaptive loci using an environment-dependent model and a population structure-based model. We assess pleiotropy by examining associations between gene expression and genetic variants. We construct gene coexpression networks to determine whether adaptive variants cluster in specific gene modules.

Our results provide new insights into the role of pleiotropy in local adaptation by examining the network properties and functional enrichment of adaptive genes in *Helianthus argophyllus*. We found that candidate locally adaptive genes identified using PCAdapt [22,23], which detects selection based solely on population structure, exhibit higher network connectivity but are not enriched in gene co-expression network modules. Adaptive loci identified through PCAdapt may function as highly connected regulatory genes whose involvement in multiple biological processes restrains their evolvability, as suggested by their low between-population genetic differentiation. In contrast, adaptive genes identified using LFMM [24], incorporating environmental associations, had lower network connectivity but were significantly enriched in specific gene network modules, suggesting a potential trade-off between pleiotropy and modular adaptation [25]. This set of candidate adaptive genes is enriched in primary metabolic pathways such as histidine biosynthesis, glycolysis, and pyruvate metabolism that appear to function within modular genetic architectures, facilitating local adaptation through specialized functional units rather than broad regulatory control. The two methods appear to identify candidate genes that likely have experienced different routes of local adaptation, possibly underlying traits that were closer to or distant from their respective fitness optima. These findings support that local adaptation involves both functional specialization (modularity) and highly pleiotropic genes acting as central hubs in gene networks [15]. This aligns with recent revisions to Fisher’s model, which propose that while pleiotropic loci can drive adaptation when phenotypes are far from their fitness optimum, modular architectures may be favored in fine-scale local adaptation, where selection acts on genes with specific metabolic or physiological roles, rather than broadly connected regulatory genes [3,4,26].

## Results

### RNA sequencing

We analyzed total RNA sequence data for 19 *Helianthus argophyllus* samples collected from North inland (late flowering, n = 9) and coastal sub-populations (early flowering, n = 10) of its range in Texas to identify genomic footprints of local adaptation. Hereafter, we refer to these populations as “north” and “coast”. A total of 6,283,383 SNPs and Indels were called from Freebayes [27]. After filtering, we retained 106,133 biallelic SNPs in 19,394 genes.

### Genetic diversity and population structure

Nucleotide diversity (*θ_π_*) and Watterson theta (*θ_W_)* had an approximate value of 0.2, indicating high levels of genetic diversity within the *H. argophyllus* population. Tajima’s *D* was slightly negative, *D* = −0.09, possibly due to the presence of rare alleles or population expansion. PCA (Fig. 1A) and *F_ST_* (mean *F_ST_* = 0.21) demonstrated genetic differentiation between the sampled sub-populations. PC1 (18.4 PVE) and PC2 (13 PVE) showed a clear differentiation between north and coast samples. Admixture analysis using NGSAdmix suggests that the population is stratified into two distinct genetic groups at K = 2, representing the two sampled populations (Fig. 1B), with differences between the populations remaining at higher K values.

**Fig 1.**
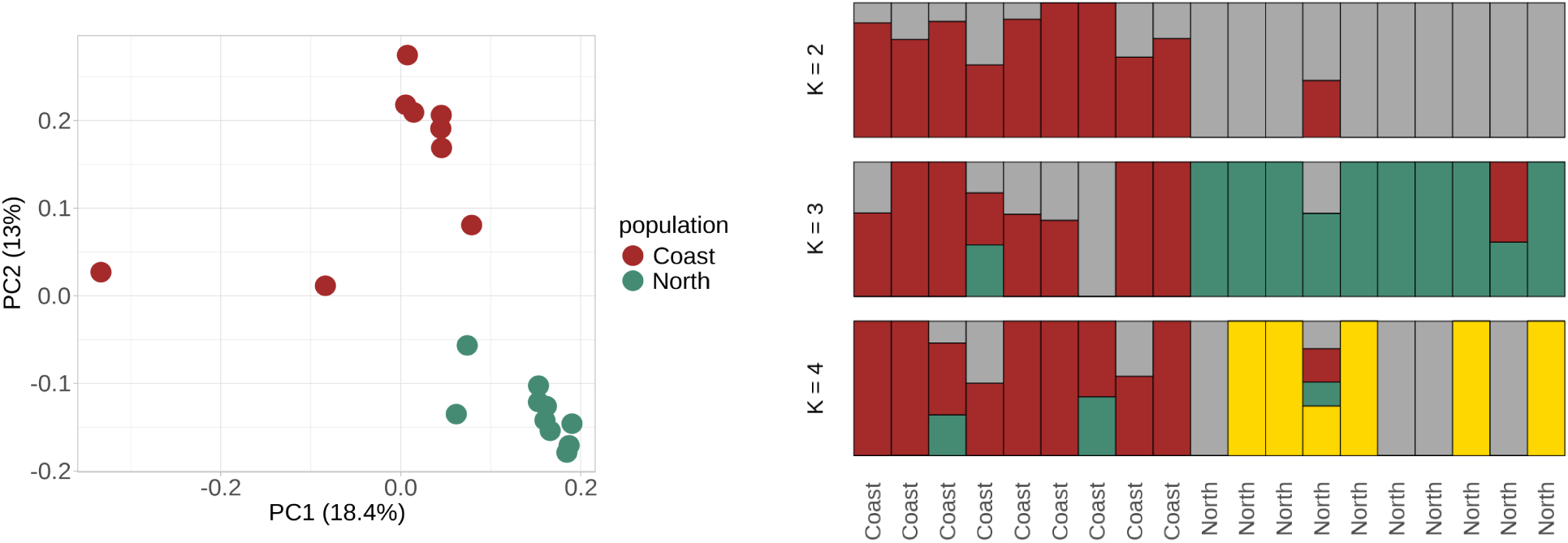
Genetic differentiation and population structure in *H. argophyllus*. **A:** Genetic differentiation along the first two PCs. Percentages in brackets show the proportion of variance explained by each PC axis. **B:** Approximate admixture proportions for each sampled individual at three different levels of K.

### Selection scans

We used two methods to identify genes underlying the local adaptation of *H. argophyllus*: LFMM 2 and PCAdapt [22,24]. LFMM 2 takes an environment-genotype association approach that combines population structure and environmental variables, while PCAdapt uses information from population structure only. For our LFMM model, we controlled for population structure using K = 4 latent factors estimated from the scree plot of the genotype-based principal component analysis (PCA) (Fig. S5). We defined the main axes of environmental variation using a principal component analysis (PCA) of 19 BIOCLIM variables. We used the first five PCs, which explain 98.95% of the total variation. Out of all 106,133 biallelic SNPs, 1,438 distributed across 560 genes bore signatures of selection in the LFMM model (*q* < 0.05). BIOCLIM PC1, PC2, PC3, PC4, and PC5 were associated with 44, 125, 1,271, 15, and 138 genes, respectively. In the PCAdapt analysis, using K = 4 latent factors, 4,968 SNPs distributed across 1,122 genes were identified as selection outliers (*q* < 0.05). Twelve SNPs in 10 genes were identified by both methods, fewer than expected by chance (*p-*value < 2.2 × 10 ^-16^, Fisher’s exact test). For further analyses, we considered genes with outlier SNPs from either approach as candidates for local adaptation (“selection outliers”). Genes containing bi-allelic SNPs not identified as selection outliers make up our control comparison group.

### Proportion of eQTLs and eGenes in selection outliers

To investigate whether selection outlier loci were more pleiotropic than control genes, we tested associations between SNPs and gene expression to identify *trans*-regulatory variants that control the expression of other genes. 42,638 genes were used for the eQTL analysis. The model included admixture proportions (K = 4) as cofactors to control for population structure. A total of 1,824 (4.3%) eQTL genes (genes containing eQTL SNPs) were identified and associated with the expression of 3,129 (7.3%) genes (eGenes) (FDR < 0.01). Twenty-six (0.06%) genes were identified as both eQTL and eGenes. Of the 560 LFMM outlier genes, 114 (20.4%) were eQTLs, while 208 (18.5%) PCAdapt outliers were eQTLs. Eleven (2%) LFMM outlier genes and 24 (2.1%) PCAdapt outlier genes were eGenes. Compared to the rest of the transcriptome, both outlier gene sets harbored more eQTLs (*p* < 2.2e-16, Fisher’s exact test) and fewer eGenes, respectively (*p* < 2.2e-16, Fisher’s exact test; Fig. 2).

**Figure 2.**
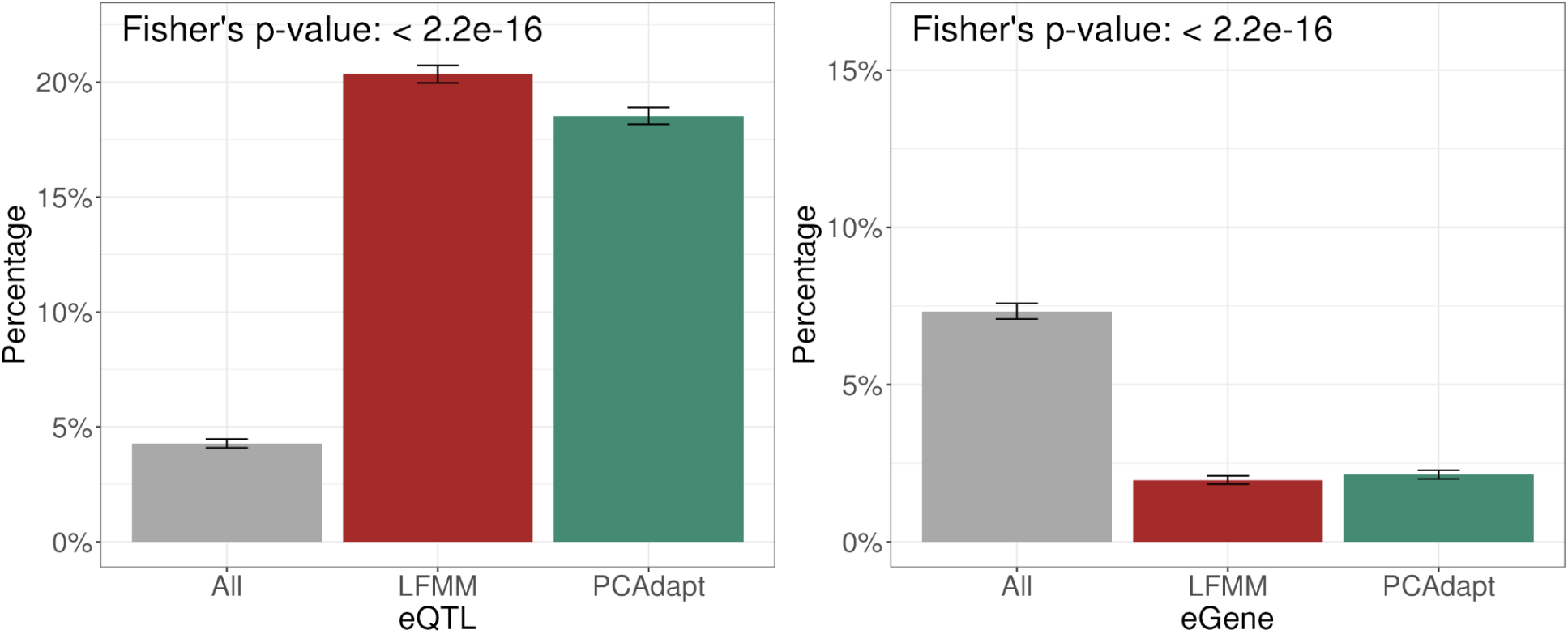
Compared to control genes, the proportion of eQTLs and eGenes in candidate adaptive genes identified by the LFMM and PCAdapt methods is significantly higher for eQTLs and lower for eGenes, respectively. Error bars were calculated using a 95% bootstrapped confidence interval.

### Evidence of selection at outlier loci

Genetic differentiation, as estimated by *FST,* was considerably higher for LFMM outlier genes than for control genes (*p*-value < 2.2e-16, Wilcoxon rank sum test). However, they were not significantly different in *D_xy_* values (Fig. 3). In contrast, PCAdapt outlier genes had lower *F_ST_* and lower *D_xy_* compared to control genes (*p-*value = 3.941e-06 and < 2.2e-16, respectively, Wilcoxon rank sum test) (Fig. 3). These results emphasize that our two selection outlier methods appear to be identifying genes with substantially different signatures of selection. We also find that *F_ST_* and *D_xy_*are significantly elevated in eQTL genes and reduced in eGenes (Fig. 3).

**Fig 3.**
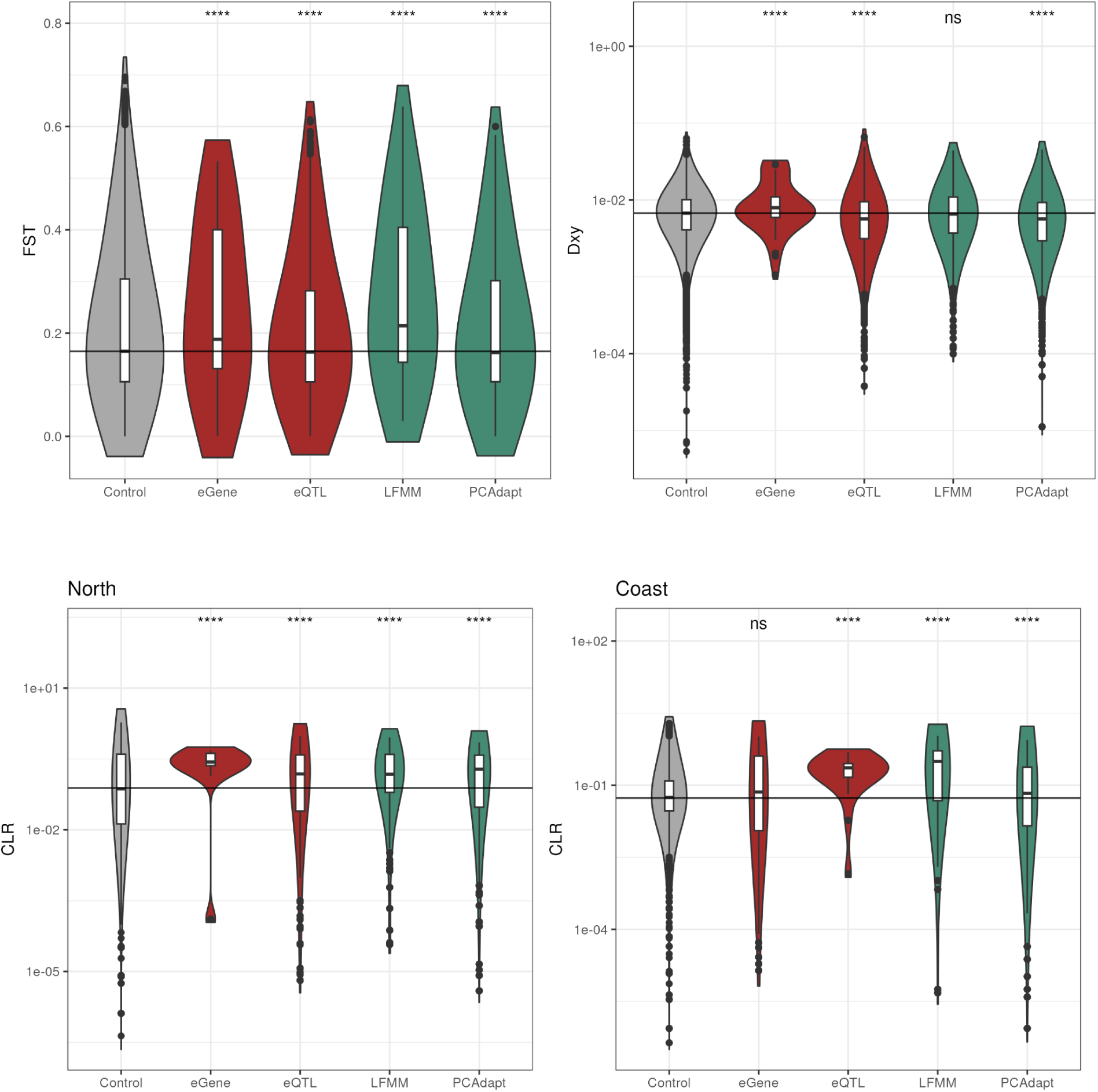
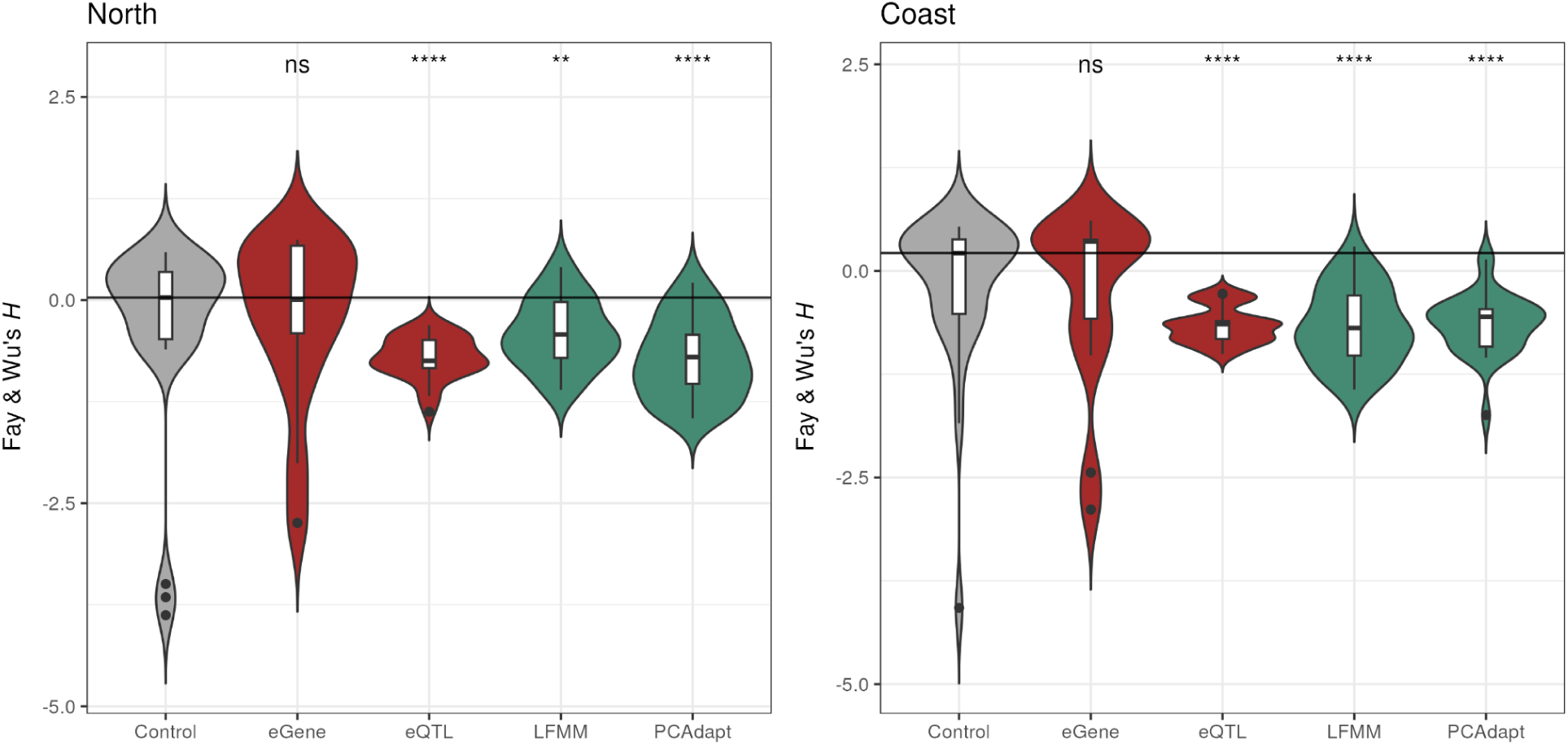
Genetic differentiation and selective sweep at adaptive genes. **A**: Weighted *F_ST_*estimates across adaptive genes, eQTLs, eGenes, and the rest of the transcriptome (control). **B**: Distribution of CLR and **C**: Fay and Wu’s *H* values. The horizontal line represents the median value of the transcriptome-wide background for all estimated values. Differences significant at p < 0.05 are indicated, even where they appear visually similar.

We estimated two measures of genetic diversity, *θ_π_* (nucleotide diversity) and *θ_W_* (Watterson’s estimator), to explore additional evolutionary patterns in outlier genes. Nucleotide diversity and Watterson’s theta should be approximately equal in a population under equilibrium conditions (evolving neutrally with constant demographic size) [28]. We would expect genes experiencing positive or purifying selection to exhibit reduced genetic diversity, while balancing selection (or a higher local mutation rate) might result in increased genetic diversity, and transcriptome-wide differences between *θ_π_* (nucleotide diversity) and *θ_W_* likely reflect population size changes. In the north population, PCAdapt outlier genes had significantly lower *θ_W_*compared to the control genes (*p-*value = 0.003819, Wilcoxon rank sum test) (Fig. S3), but *θ_π_* was not significantly different (*p*-value = 0.3752, Wilcoxon rank sum test). A similar pattern was observed for PCAdapt outliers in the coast population, although *θπ* was significantly larger than in the control (*θ_π_*; *p-*value = 0.004505, *θ_W_*; *p-*value = 0.001016, Wilcoxon rank sum test). In the north population for LFMM outlier genes, *θπ* was significantly higher than the control, while *θ_W_* was not significantly different (*θ_π_*; *p-*value *<* 2.2e-16, *θ_W_*; *p-*value = 0.06571, Wilcoxon rank sum test). An opposite pattern was observed in the coast population: LFMM outlier genes exhibited significantly lower *θ_W_* compared to the control (*p*-value *=* 1.57e-14, Wilcoxon rank sum test) and no significant difference in *θ_π_* (*p-*value = 0.2015, Wilcoxon rank sum test).

Finally, we further validated our outlier genes using Fay and Wu’s *H* neutrality measure and the Composite Likelihood Ratio (CLR) test for selective sweeps (Fig. 2B). CLR estimates at the candidate adaptive genes identified by the LFMM and PCAdapt methods, as well as eQTL and eGenes, were significantly more positive compared to the control genes (*p*-value < 2.2 × 10 ^-16^, Wilcoxon rank sum test for both LFMM and PCAdapt) (Fig. 2B). PCAdapt and LFMM outliers in both populations had more negative Fay & Wu’s *H* estimates compared to control genes (LFMM-north; *p-*value = 0.001; PCAdapt-north; *p-*value = 6.623e-06; LFMM-coast; *p-*value = 4.998e-05, PCAdapt-coast; *p-*value = 5.977e-05) (Fig. S4). These results suggest that both candidate locally adapted gene sets are enriched for selective sweeps in both populations.

### Connectivity of selection outliers

We estimated gene co-expression networks to further evaluate the pleiotropy of genes harboring selection outliers. Our gene co-expression analysis identified 44 modules, with a median of 41 genes per module (range, 9–2,209). Three modules contained more than expected LFMM genes (*p*-value < 0.05, Fisher’s exact test), while no WGCNA module had significant enrichment for PCAdapt genes. We used the connectivity of our candidate adaptive genes, as well as our eQTLs and eGenes, to estimate transcriptomic pleiotropy. More highly connected genes are putatively more pleiotropic in that changes to them will have broader effects on the expression of other genes. PCAdapt outlier genes and eQTLs had higher connectivity than control genes (*p*-value < 2.2 × 10 ^-16^, Wilcoxon rank sum test). In contrast, LFMM outlier genes and eGenes were less connected than the control (*p*-value < 2.2 × 10 ^-16^, Wilcoxon rank sum test) (Fig. 4).

**Fig 4.**
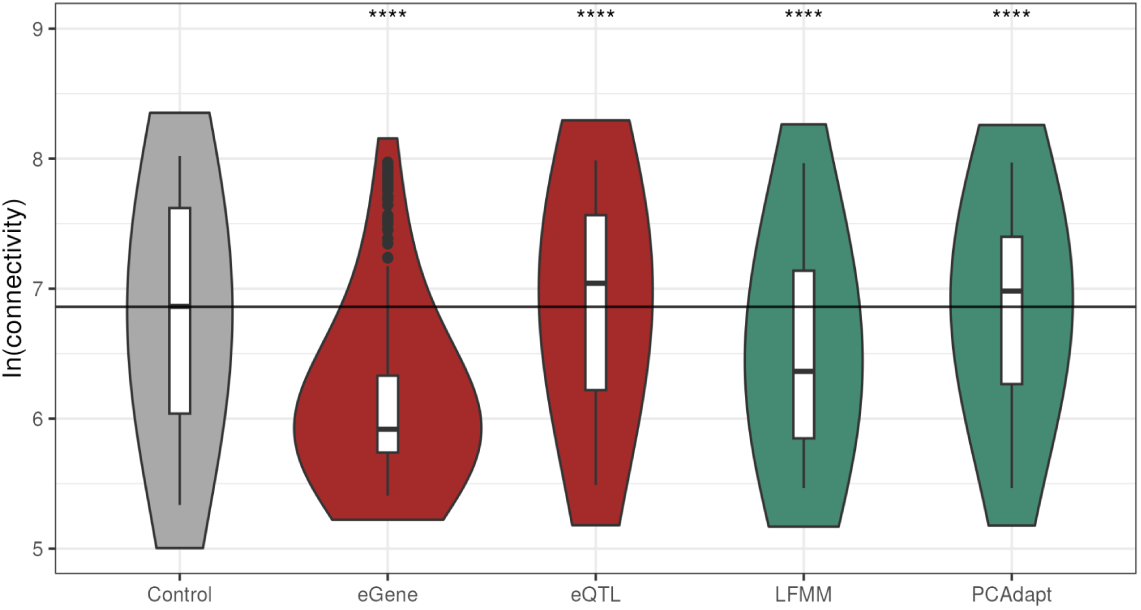
Gene network connectivity at candidate adaptive genes. Estimated connectivity of candidate adaptive genes, eGenes, and eQTLs compared to the transcriptome-wide control.

### Annotation of adaptation loci and gene co-expression networks

Our gene overrepresentation analysis revealed significant enrichment for pollen recognition and protein serine/threonine kinase activity in both LFMM and PCAdapt candidate genes (Fig. 5). LFMM genes were additionally enriched in functions related to sulfotransferase, catalase, and oxidoreductase activities. In contrast, PCAdapt genes were significantly enriched for signal transduction, defense response, and dehydrogenase activity. The few genes identified by both methods included members of the bZIP family of transcription factors, whose functional orthologues in *Arabidopsis thaliana* show high expression in reproductive tissues such as pollen, inflorescence meristem, carpels, and petals. Several of these genes also exhibited high expression in guard cells, the shoot apex, and the shoot system [29]. WGCNA modules with more than expected LFMM candidate genes were involved in pathways annotated as predominantly primary metabolism, including histidine biosynthesis, de novo purine biosynthesis, pyruvate oxidation, triacylglycerol biosynthesis, and leucine biosynthesis (Table 1).

**Fig 5.**
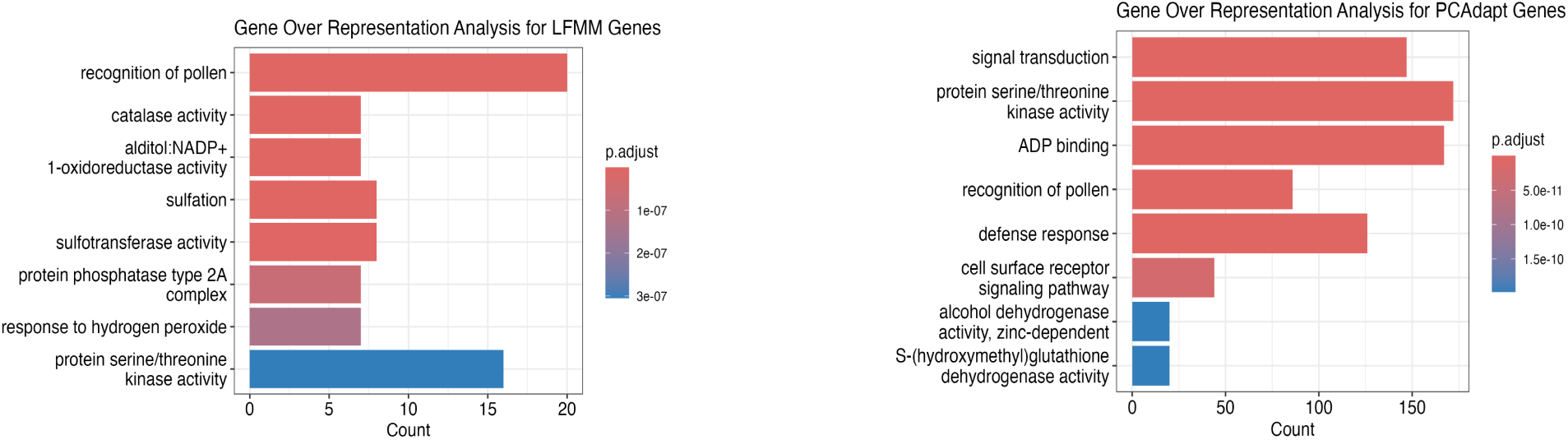
Gene ontology enrichment for biological processes of adaptive genes identified by LFMM (top), PCAdapt (middle), and both LFMM and PCAdapt (bottom).

**Table 1.** KEGG Module Annotation of the three network modules containing more than expected LFMM adaptive genes.

| WGCNA Module | KEGG ID | Description | Adjusted p-value |
| --- | --- | --- | --- |
| Dark grey | M00026 | Histidine biosynthesis, PRPP => histidine | 0.00063984 |
|  | M00129 | Ascorbate biosynthesis, glucose-1P => ascorbate | 0.0014631 |
| Dark turquoise | M00002 | Glycolysis, a core module involving three-carbon compounds | 0.0006257 |
|  | M00048 | De novo purine biosynthesis, PRPP + glutamine => IMP | 0.00117279 |
|  | M00001 | Glycolysis (Embden-Meyerhof pathway), glucose => pyruvate | 0.00228412 |
|  | M00130 | Inositol phosphate metabolism, PI=> PIP2 => Ins(1,4,5)P3 => Ins(1,3,4,5)P4 | 0.00288918 |
| **Green** | M00307 | Pyruvate oxidation, pyruvate => acetyl-CoA | 0.00107016 |
|  | M00089 | Triacylglycerol biosynthesis | 0.00954748 |
|  | M00432 | Leucine biosynthesis, 2-oxo isovalerate => 2-oxo isocaproate | 0.03465363 |

## Discussion

Pleiotropy is a key factor in adaptive evolution, influencing how genetic loci contribute to phenotypic variation and shaping evolutionary trajectories [1,2]. Fisher’s geometric model posits that adaptation is more efficient through small-effect loci, as large-effect mutations may overshoot the fitness optimum and cause maladaptation [3]. However, Orr’s extension of Fisher’s model suggests that pleiotropic loci can facilitate rapid adaptation in novel environments, particularly following range expansion [7,30]. The role of pleiotropy in adaptation has been debated, with some models suggesting that highly pleiotropic loci constrain evolution by generating trade-offs [25,31]. In contrast, others propose that such loci can accelerate adaptation [32], particularly when populations are far from optimal fitness [4,33].

Our findings provide new insights into the interplay between pleiotropy and local adaptation in the silverleaf sunflower *Helianthus argophyllus*. We identified distinct sets of candidate adaptive loci using two methods: PCAdapt and LFMM selection scans. Both sets are enriched for loci associated with the expression of other genes (eQTL) and signals of positive selection. PCAdapt outliers, identified through controlling for population structure, exhibited significantly higher gene co-expression network connectivity than the transcriptome-wide control without specific enrichment in any co-expression module. This suggests that the candidate adaptive loci identified by PCAdapt fit Orr’s extension of Fisher’s model and represent regulatory hub genes with broad pleiotropic effects, influencing multiple biological processes but potentially facing evolutionary constraints due to their central role in gene networks.

In contrast, LFMM outliers, identified by incorporating environmental variables, had significantly lower gene network connectivity and were enriched in gene co-expression modules involved in primary metabolic pathways such as histidine biosynthesis, glycolysis, and pyruvate metabolism. This suggests these candidate genes may contribute to adaptation through modular genetic architectures rather than extensive pleiotropy [34]. These two distinct sets demonstrate that adaptation may take multiple avenues, even within the same populations of the same species.

Our results are consistent with those of Lotterhos *et al.* (2018), who examined how genetic architecture influences local adaptation to climate in lodgepole pine (*Pinus contorta*) [35]. Their results support a modular pleiotropy hypothesis, suggesting that candidate locally selected genes play a significant role in specific environmental conditions but have little effect across environments [3,15]. For example, a gene might regulate multiple traits related to temperature adaptation (e.g., cold tolerance and seasonal growth timing) but have little or no effect on drought response or nutrient uptake. This modular genetic structure suggests that adaptation can be specialized, with different genes fine-tuned for particular environmental challenges rather than broadly shaping responses. Such modular architectures may be favored in complex environments where multiple selective pressures, such as temperature, precipitation, and soil composition, act independently. This makes it more advantageous for genes to affect specific traits peculiar to a particular environmental context than to influence many unrelated factors universally [15, 16].

The higher proportion of eQTLs in candidate adaptive genes compared to non-adaptive genes suggests that regulatory variation plays a significant role in the local adaptation of *Helianthus argophyllus*. This could be because adaptive genes are subject to more plastic gene expression control, potentially enhancing their ability to respond to environmental selection [36].

Interestingly, while PCAdapt outliers exhibited higher network connectivity, they were not significantly enriched in any co-expression modules. This suggests that although they function as regulatory hubs, they may not form distinct adaptive modules, a pattern potentially reflecting evolutionary constraints imposed by widespread pleiotropy [25]. In contrast, LFMM outliers were more likely to be involved in specific co-expression modules, reinforcing that environmental selection may act through functionally specialized genetic networks.

Our results also provide insights into the selective pressures acting on these candidate adaptive loci. Genetic differentiation (*F_ST_*) was significantly higher for LFMM outliers than for PCAdapt and control genes. Still, these loci did not show significant differences in *D_xy_* values from the control, suggesting that they may be undergoing selection within locally adapted populations without widespread divergence [38]. This pattern is consistent with a modular adaptation model, where environmental selection acts on functionally specialized loci without extensive genetic differentiation [15,35]. PCAdapt outliers exhibited lower *F_ST_* and *D_xy_* than the control, suggesting that these loci may be subject to purifying or balancing selection [15,35].

The patterns of diversity observed between PCAdapt and LFMM outliers in both populations suggest the different ways selection may act on these loci. The elevated *θ_π_* and lower *θ_W_* and the strong sweep signals observed in CLR and Fay & Wu’s *H* in LFMM outliers in the north population suggest that selection may be acting on standing genetic variation. In such cases, a hard sweep can still arise if one beneficial allele confers a stronger advantage or if demographic constraints reduce haplotypic diversity, causing selection to favor a single variant [39, 40]. The coast population, on the other hand, shows reduced diversity at LFMM loci, indicative of a hard sweep where a new advantageous mutation has rapidly fixed. These contrasting patterns suggest that while both populations might adapt to similar selective pressures, the source and dynamics of the beneficial alleles differ, possibly due to variations in demographic history, gene flow, or the timing of selection events.

Gene ontology analysis revealed functional differences between adaptive loci identified by LFMM and PCAdapt. LFMM outliers were significantly enriched in primary metabolic pathways, suggesting that modular adaptation may be driven by genes involved in biochemical processes [38]. In contrast, PCAdapt outliers were enriched in pathways related to broader regulatory functions, such as signal transduction, defense response, and protein serine/threonine kinase activity. This is consistent with the role of pleiotropic regulatory genes in mediating the evolution of complex traits.

The distinction between the adaptive loci identified by PCAdapt and LFMM underscores a potential trade-off between pleiotropy and modularity in local adaptation. Due to their broad regulatory functions, highly pleiotropic loci, such as those identified by PCAdapt, may play a role in long-term evolutionary processes. In contrast, adaptive loci functioning within modular pathways may allow for more flexible and rapid responses to environmental selection. This is consistent with recent extensions of Fisher’s geometric model, which propose that while pleiotropic loci can facilitate adaptation in populations far from their fitness optimum, modular architectures may be more advantageous for fine-scale adaptation, where selection acts on genes with specific metabolic or physiological roles, rather than broad regulatory influence [7,18].

While our study provides new insight into *H. argophyllus* adaptive variation, some limitations are to be considered. First, our transcriptomic data were collected at a single developmental time point (3 weeks), and our analyses might only show how *H. argophyllus* adapts during this growth period and would not reflect the entire trajectory of *H. argophyllus* adaptation. Secondly, the differences we observe might be due to neutral processes, since our analysis approach only captured relevant genomic variation within genes.

Our study shows how important it is to think about both pleiotropy and modularity when studying local adaptation. Highly pleiotropic genes can act as central hubs, which may limit a species’ ability to evolve. In contrast, modular genetic systems allow for adaptation by focusing on specific functions. Backed by genetic differentiation data, eQTL analysis, selective sweep signatures, and gene ontology enrichment, our findings demonstrate how gene networks and specialized functions shape local adaptation in *Helianthus argophyllus*. This research contributes to the ongoing discussion of how selection, pleiotropy, and modularity work together to drive evolution in complex organisms.

## Conclusions

This study provides new insights into the relationship between pleiotropy and adaptation in a naturally diverging plant species. Our findings suggest that candidate locally adaptive loci in *Helianthus argophyllus* are under strong selection and more likely to regulate gene expression than non-adaptive loci. Our results also suggest the need to use multiple approaches when identifying potential loci under selection, as each method may identify unique variations. In our case, one method identified genes that are highly connected within gene co-expression networks, as well as other genes at the periphery of these networks. Local adaptation may be facilitated and hindered by pleiotropy, which likely depends on the distance of the populations from fitness optima and the consistency of those optima. Understanding this dynamic helps us understand how genetic structures drive adaptation, with important implications for evolutionary biology, conservation, and plant breeding.

## Acknowledgments

We thank Dr. Andrew Whitehead (UC Davis) for his valuable insights on trait evolvability. We are also grateful to the members of the Moyers Lab at UMass Boston for their support and contributions. The Unity Cluster hosted by UMass Amherst was used for all computational analyses.

## Material and methods

### Sample collection and RNA sequencing

Sample collection and sequencing are described in Renaut *et al*. 2013. Here, we include 19 populations representing two distinct subpopulations of *H. argophyllus,* identified by previous research [41]. Briefly, individual seeds representing unique populations of *H. argophyllus* collected by BTM in 2011 were grown for 3 weeks in a growth chamber with 12 hours of daylight at 22 °C [41]. All above-ground tissue was then frozen and stored at −80° until RNA was extracted using a modified TRIzol Reagent protocol. Libraries were prepared using mRNASeq (Illumina, San Diego, CA) approach and sequenced on a GAII (paired-end sequencing, 2 × 100 bp reads). The average number of reads per individual was 1864375, with an average read depth of 17.8.

### Sequence processing

We ran FastQC/0.11.5 [42] on the RNA sequencing reads to assess quality and Trimmomatic/0.39 [43] to perform sliding window trimming with a 4-base window size. We removed reads with an average quality below 20 and reads shorter than 25 bases after trimming. We aligned reads to the Ha412HOv2 *H. annuus* reference genome (https://www.sunflowergenome.org/assembly-data/) [44] using STAR [45]. After alignment, we used Picard/2.23.3 [46] to add read group information and remove duplicated reads. We called variants on aligned reads using Freebayes/1.3.5 [27] and filtered the VCF file with vcftools 0.1.14 [47]. We applied the following filtering criteria: site quality ≥ 30, genotype quality ≥ 20, read coverage ≥ 6, and missing data < 20%.

### Genetic diversity and population structure

We filtered genotype variants for minor allele frequency (MAF) ≥ 0.05 for these analyses. We estimated *θ_W_*, *θ_π_*, Tajima’s *D*, and *F_ST_* with ANGSD/0.935 [48] and used Pixy to estimate *Dxy* for all sampled individuals [49], and used NGSAdmix to identify admixture proportions [50]. We estimated θ_W_, θ_π_, and Tajima’s *D* within each population. Waterson’s θ (*θ_W_*) is an estimator of population mutation rate and can be compared to nucleotide diversity (*π or θ_π_*), a measure of within-population polymorphism, to identify genetic loci that have experienced non-neutral evolution or demographic changes (Watterson 1975, Tajima 1989). We estimated mean *F_ST_* within 5000 kb sliding windows. We used the *H. annuus* reference as an outgroup when calculating Fay and Wu’s *H* to estimate the proportion of derived alleles.

### Selection outliers

We used the LFMM 2 and PCAdapt R packages to identify selection outliers at a *q*-value threshold of <0.05. LFMM 2 is a genotype-environment association (GEA) method that incorporates more than one independent variable in a multivariate model. These variables can be climatic, methylation, or gene expression data [24]. We included 19 BIOCLIM variables in our LFMM 2 model to identify genotype-environment associations [51]. PCAdapt identifies locally adapted candidates after controlling for population structure. PCAdapt is optimized to handle admixed individuals, account for population divergence, handle large sequencing datasets, and has a relatively low FDR [22].

### eQTL and eGenes

We counted the number of reads expressed for each gene using FeatureCounts in the Subread/1.6.2 package [52]. We normalized the gene count file generated by Subread for our eQTL analysis with the Deseq2 R package [53], which uses the median-ratio normalization method [54]. We used the MatrixEqtl R package to identify loci (eQTL) associated with changes in the expression of specific genes (eGenes) [55]. We increased our filter MAF threshold from 0.05 to 0.1 to reduce the occurrence of spurious associations [56]. To control for expression heterogeneity, we performed a principal component analysis on our expression data following [57] and selected the first two principal components that explained 92% of the variation in our expression data. We included the residuals from the regression analysis using the first two principal components in our eQTL model to control for the effect of population structure [57].

### Selective sweeps

We subset the VCF file by the genomic locations of selection outlier SNPs or genes, eQTL, eGenes, and control genes for each population. Control genes were the expressed genes that were not identified by any of the other analyses. We identified putative selective sweeps using two approaches: Fay and Wu’s *H* [58] and the Composite Likelihood Ratio (CLR) method using the SweepFinder2 package [59] (Vy and Kim, 2015). In estimating CLR, we subset each VCF file by chromosome and used all positions per chromosome to create grid files for the analysis.

### Statistical comparisons

We compared values for population genetic estimators across selection outlier genes, eQTL, eGenes, and control genes using the Wilcoxon signed-rank test in R [60].

### Gene coexpression networks

We used the WGCNA R package to construct gene coexpression networks [61]. We set a power threshold of six to get the adjacency matrix of the signed network using the ‘softConnectivity’ function. This helps to estimate the number of genes connected to each node in a network module [61,62]. We compared the log-transformed connectivity of node genes that were selection outliers, eQTLs, or eGenes to the connectivity of node genes in the transcriptome-wide background using the Wilcoxon signed-rank test [60]. We visualized gene coexpression modules using Cytoscape [63].

### Annotation of genes and network modules

We used the clusterProfiler R package for gene and KEGG module overrepresentation analyses of selection outlier genes and coexpression network modules [64]. Since VCF was aligned to the HA412-HO reference genome, but the HanXRQr2.0-SUNRISE genome is used in the KEGG database, we also aligned our transcriptome to the HanXRQr2.0-SUNRISE genome to extract NCBI gene IDs for each gene sequence using the NCBI BLAST command line tool [65]. We then used these gene IDs as queries in the gene overrepresentation analysis using the ‘enricher’ function in the clusterProfiler package. We used gene IDs as queries for the KEGG module analysis using the ‘enrichMKEGG’ function in clusterProfiler. All scripts can be accessed here: https://github.com/Uzezi93/Pleiotropy-and-Local-Adaptation-Silverleaf-Sunflower.

## Supporting information

**S1 Fig.**
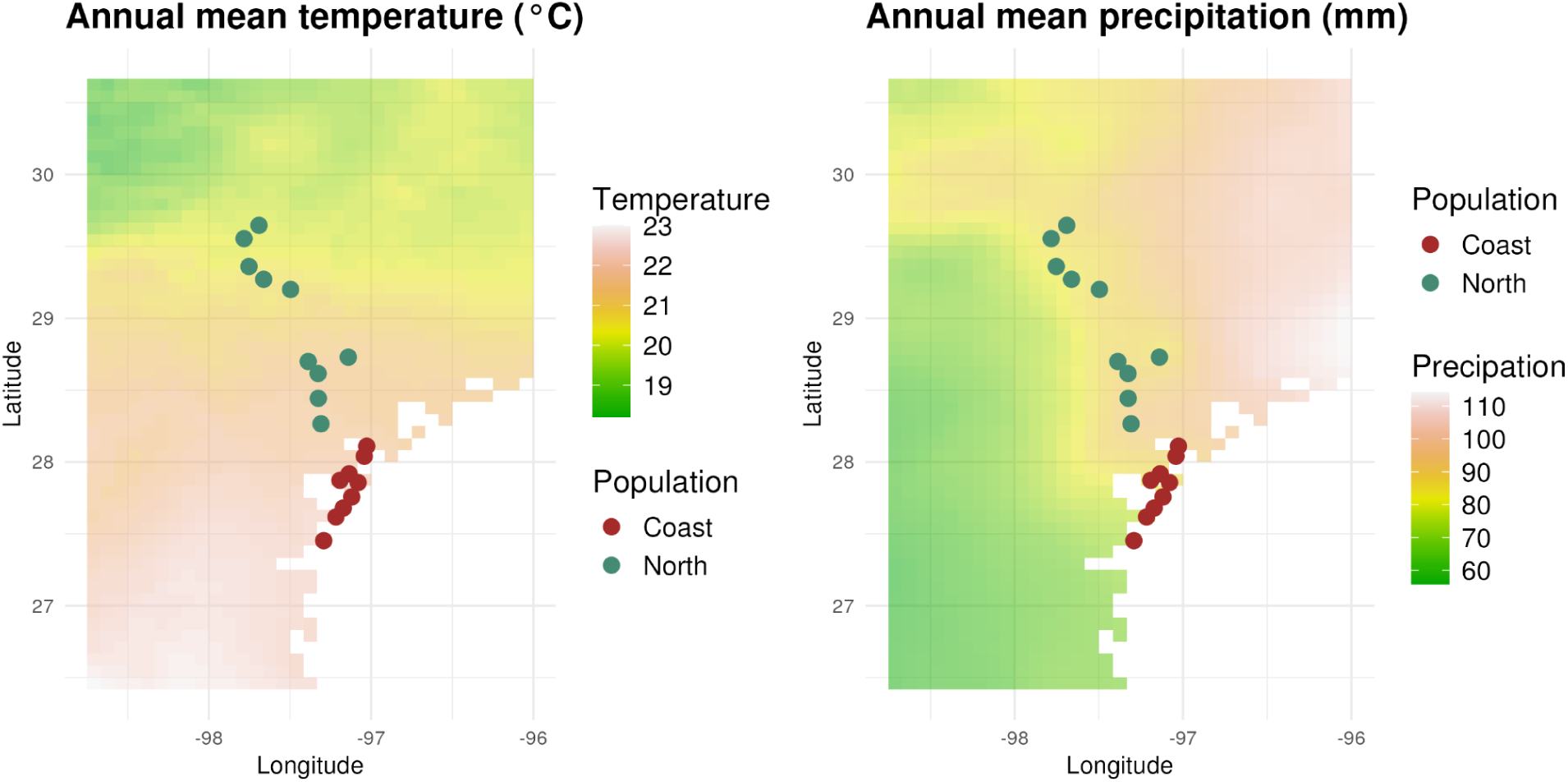
**Variation in temperature and precipitation in *H. argophyllus* sampling distribution**

**S2 Fig.**
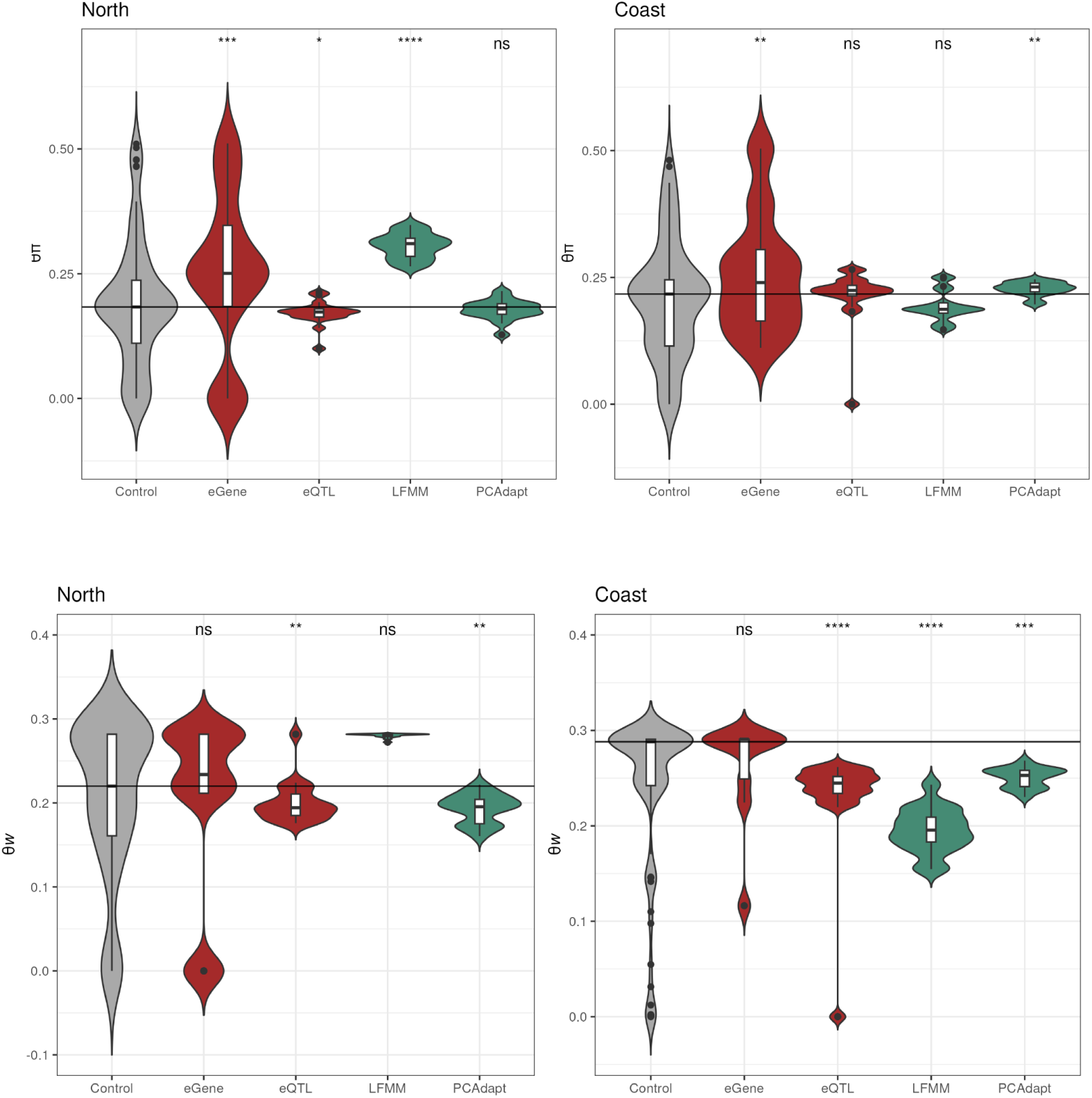
Genetic diversity in selection outliers in North and Coast Populations

**S3 Fig.**
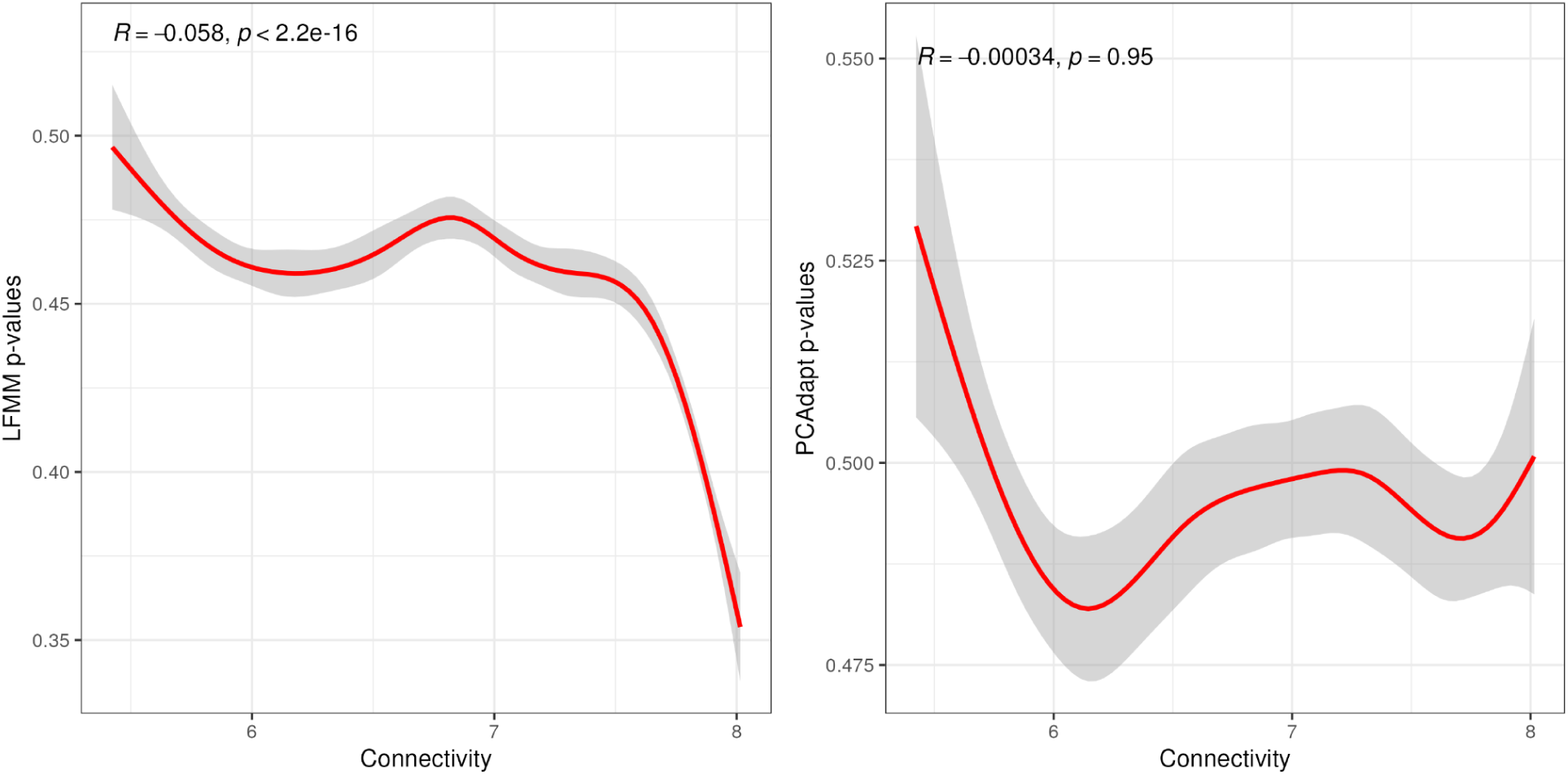
**Association between LFMM and PCAdapt connectivity and *P*-values at 95% CI**

**S4 Fig.**
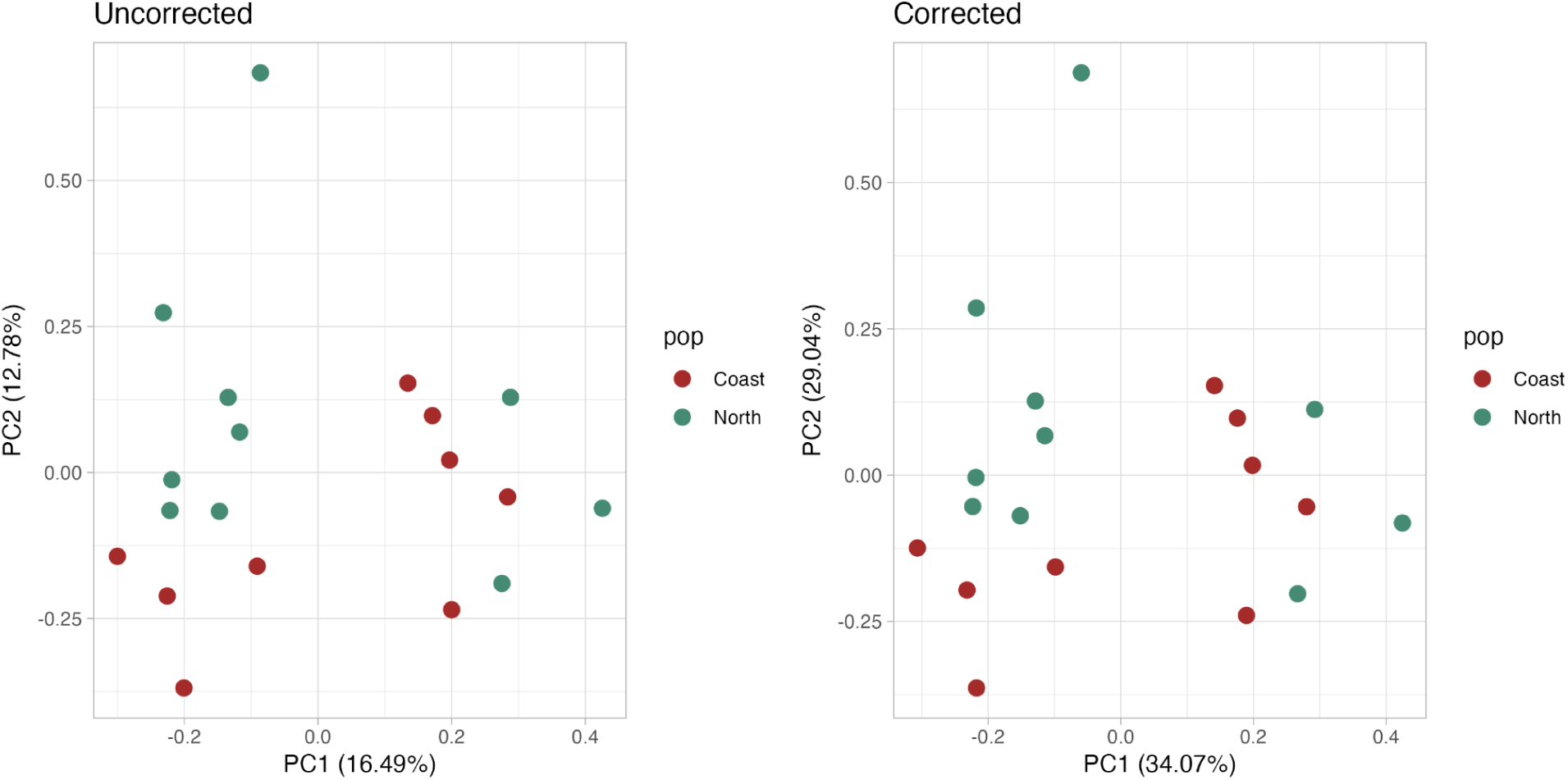
**Corrected and Uncorrected Expression-level PCA with first two PCs.**

**S5 Fig.**
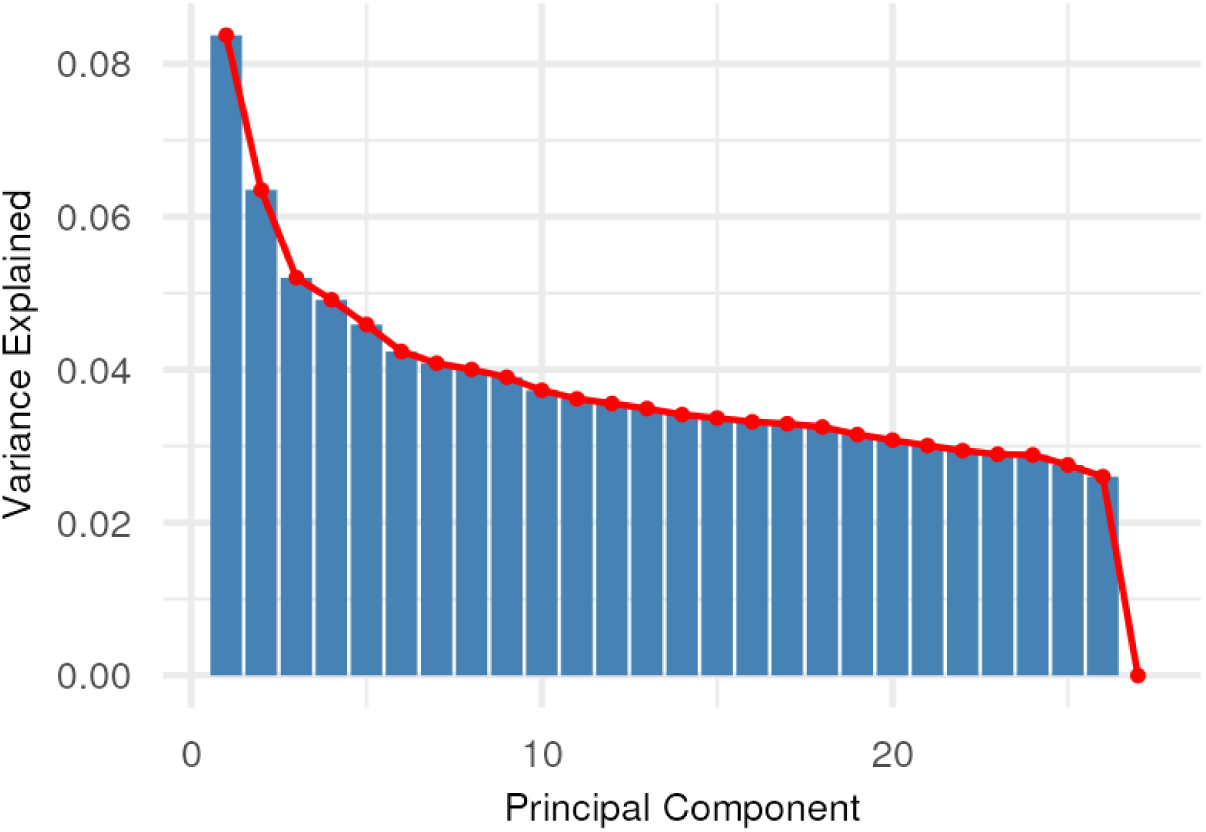
**Screeplot from genotype-based PCA**

**S6 Fig.**
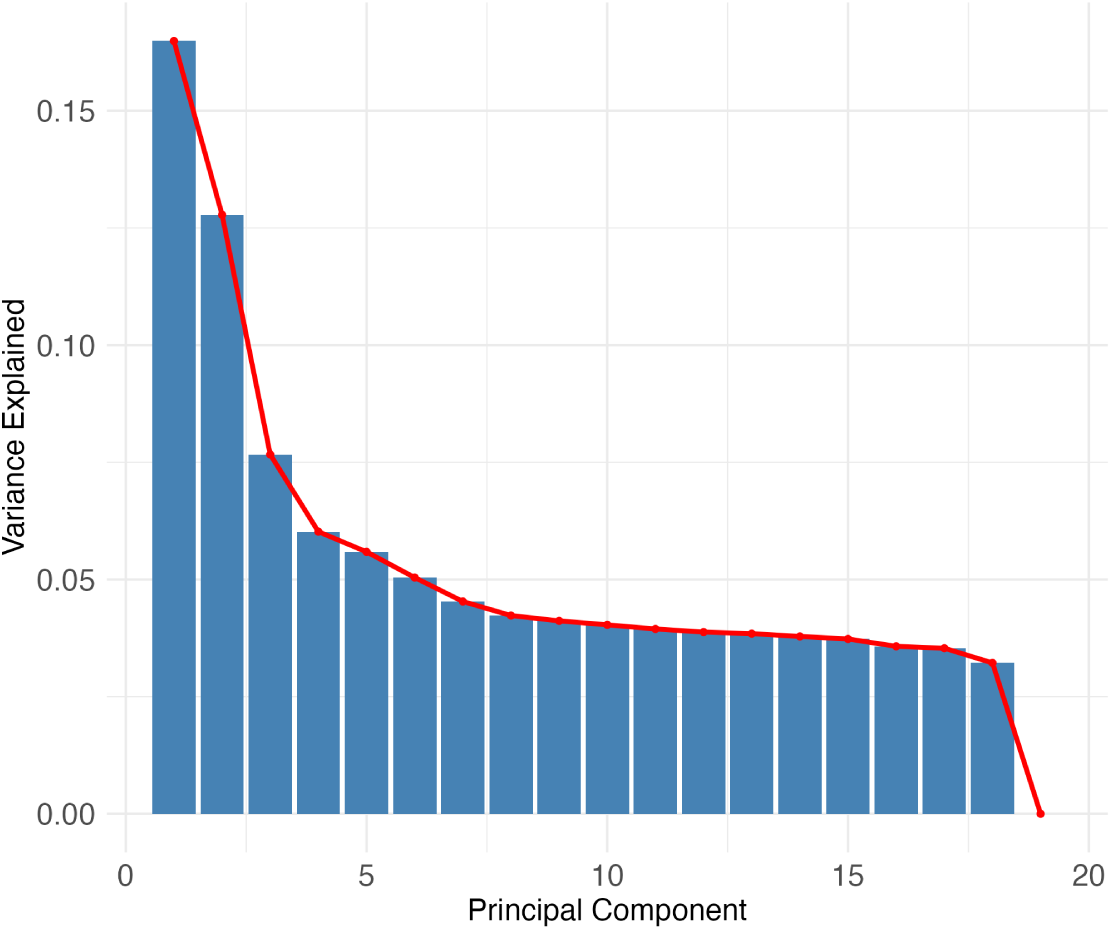
**Screeplot from expression-based PCA**

**S1 Table.** Mean estimates of genetic diversity across all expressed genes in sampled populations.

| | $\theta\pi$ | $\theta w$ | Tajima's $D$ |
| --- | --- | --- | --- |
| **North** | 0.1937444 | 0.2076168 | -0.1745136 |
| **Coast** | 0.2060006 | 0.2353111 | -0.2985429 |

**S2 Table.**
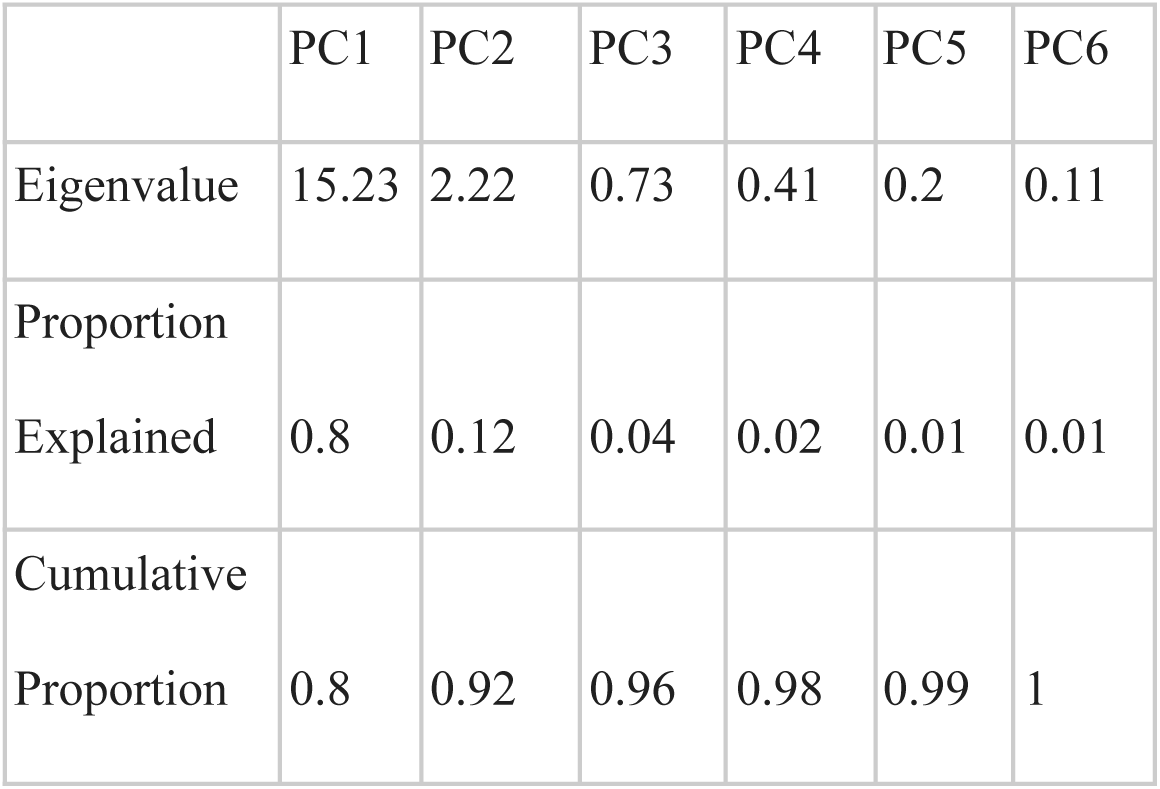
Proportion of Variance Explained (PVE) by the first six Principal Components (PC)

|  | PC1 | PC2 | PC3 | PC4 | PC5 | PC6 |
| --- | --- | --- | --- | --- | --- | --- |
| Eigenvalue | 15.23 | 2.22 | 0.73 | 0.41 | 0.2 | 0.11 |
| Proportion Explained | 0.8 | 0.12 | 0.04 | 0.02 | 0.01 | 0.01 |
| Cumulative Proportion | 0.8 | 0.92 | 0.96 | 0.98 | 0.99 | 1 |

